# Isolation of wild yeasts from Olympic National Park and *Moniliella megachiliensis* ONP131 physiological characterization for beer fermentation

**DOI:** 10.1101/2021.07.21.453216

**Authors:** Renan Eugênio Araujo Piraine, David Gerald Nickens, David J. Sun, Fábio Pereira Leivas Leite, Matthew L. Bochman

## Abstract

Thousands of yeasts have the potential for industrial application, though many were initially considered contaminants in the beer industry. However, these organisms are currently considered important components in beers because they contribute new flavors. Non-*Saccharomyces* wild yeasts can be important tools in the development of new products, and the objective of this work was to obtain and characterize novel yeast isolates for their ability to produce beer. Wild yeasts were isolated from environmental samples from Olympic National Park and analyzed for their ability to ferment malt extract medium and beer wort. Six different strains were isolated, of which *Moniliella megachiliensis* ONP131 displayed the highest levels of attenuation during fermentations. We found that *M. megachiliensis* could be propagated in common yeast media, tolerated incubation temperatures of 37°C and a pH of 2.5, and was able to grow in media containing maltose as the sole carbon source. Yeast cultivation was considerably impacted (*p*<0.05) by lactic acid, ethanol, and high concentrations of maltose, but ONP131 was tolerant to high salinity and hop acid concentrations. This is one of the first physiological characterizations of *M. megachiliensis*, which has potential for the production of beer and other fermented beverages.

## 1. Introduction

Beer is produced using four main ingredients: malted grains (usually barley), water, hops, and yeast (mostly *Saccharomyces pastorianus* and *S. cerevisiae*) (Bokulich and Bamforth, 2013). Searching for new products, additional ingredients have been introduced to beer recipes, such as fruits, herbs, and wood. Additionally, brewers are exploring new combinations of traditional ingredients, for example the quantity and varieties of hops, malting different grains or the use of non-conventional/wild yeasts during fermentation (Donadini and Porretta, 2017). It is estimated that thousands of these strains, if not species, of wild yeasts have potential for industrial application (Gutiérrez et al., 2018). Among them, some species in the *Brettanomyces, Candida*, and *Pichia* genera were initially considered only as contaminants of beer and wine (Campbell, 2003). However they are currently considered important components in high value-added beers, as they can produce sought-after flavors and aromas during fermentation (Michel et al., 2016).

Yeasts are ubiquitous in the environment and are often isolated from sugar-rich sources, such as fruit, berries, and plant exudates (Rao et al., 2008; Tikka et al., 2013), with soil and some insects being natural reservoirs of fermentative yeasts (Barry et al., 2018; Osburn et al., 2016). Multiple techniques for the isolation of wild yeasts are already well stablished. However, the process becomes complex when there is a need to characterize these isolates before their application in fermentation processes. Various tests must be performed to determine characteristics such as alcohols tolerance, ability to metabolize different types and concentrations of carbohydrates, and survival in adverse conditions (e.g., pH, temperature) (Tikka et al., 2013). In the screening process for yeasts with potential in brewing industry, it is of great importance to identify the aromatic compounds produced during and after fermentation (e.g., esters, fusel alcohols, phenols), their flocculation and attenuation profiles, and especially their growth characteristics (Osburn et al., 2016). As with *S. cerevisiae*, their domestication becomes important, selecting and maintaining wild species to obtain variants that are able to develop in a controlled way, even under suboptimal conditions compared to those observed in their natural environment (Gallone et al., 2016; Steensels et al., 2019).

Thus, the objective of this work was to isolate wild yeasts from their natural habitat, selecting them for their application potential for beer production and physiologically characterizing them, aiming at the future domestication and establishment of the cultures as commercial starters for the brewing market. Here, one such strain, *Moniliella megachiliensis* ONP131, showed great promise for beer production. Wild yeasts of the *Moniliella* genus, which is still under studied regarding its taxonomy and ecology. This yeasts in this genus are mainly found in tropical ecosystems, present in flowers (Thoa et al., 2015), unprocessed foods, and insects (Lachance et al., 2001; Rosa et al., 2009). Currently, the biotechnological importance of these yeasts lies in the commercial production of erythritol, a natural four-carbon sugar alcohol that is a non-carcinogenic sweetener (Thoa et al., 2015). *Moniliella megachiliensis* has been reported for its large erythritol production capacity (Thoa et al., 2015), including patent-protected processes (Ghislain et al., 2002). It has further been described that this yeast is able to metabolize large concentrations of glucose and maltose (Hoog et al., 2011), which are the main carbohydrates found in beer worts. Although used extensively for erythritol production, there are few studies regarding its application in other fermentation processes. We found that *Moniliella megachiliensis* ONP131 could be propagated using traditional yeast growth media, tolerated various stressors associated with beer fermentation, and produced beer with a pleasant organoleptic profile.

## 2. Material and Methods

### 2.1. Wild yeast isolation and identification

Fifteen samples from sources such as soil, tree bark, roots, leaves, and flowers were collected in Olympic National Park (Port Angeles, Washington, US) in August, 2019. Samples were stored in plastic zip type bags and stored at 4°C until processing. A small piece of 2 cm^2^ of each sample was aseptically extracted and added to 5 mL of YPM8E5 medium (10 g/L yeast extract, 20 g/L peptone, 80 g/L maltose, 5% ethanol (v/v), 50 µg/mL kanamycin, 50 µg/mL chloramphenicol), and then incubated with shacking for 72 h at 30°C. A volume of 10 µL of each sample was used for microbial isolation on Wallerstein Laboratory Nutrient agar (WLN) medium, and plates were incubated as described above. Colonies with yeast morphology were plated again on YPD agar medium (10 g/L yeast extract, 20 g/L peptone, 20 g/L dextrose and 20 g/L agar), inoculated into 10 mL of YPD medium and then visualized by phase contrast microscope at 100x magnification to determine morphology and purity. Saturated cultures of pure isolates were mixed with 30% glycerol and stored at −80°C to establish a culture bank.

For species identification, one colony of each isolate was cultivated in liquid YPD medium, and then the cell pellet from 200 µL of culture was used for genomic DNA (gDNA) extraction. Subsequently, 0.5 µL of gDNA was used as the template for PCR with primers NL1 (5’-GCATATCAATAAGCGGAGGAAAAG-3’) and NL4 (5’-GGTCCGTGTTTCAAGACGG-3’) to amplify the variable domain (D1/D2) of the 26S rRNA gene, yielding PCR products of ∼600 bp. PCRs were performed in an Eppendorf Mastercycler pro S thermocycler following the protocol: initial denaturation at 98°C for 5 min; 30 cycles of denaturation at 98°C for 30 s, annealing at 55°C for 30 s, and extension at 72°C for 1 min; and ending with a final extension at 72°C for 10 min. PCR results were visualized using 1% agarose gel electrophoresis for 1 h at 130 V.

Amplified fragments were purified using a PCR Purification kit (ThermoScientific, Walthan, MA), quantified using a Nanodrop spectrophotometer (ThermoScientific, Walthan, MA), and submitted for Sanger sequencing by ACGT Inc (Wheeling, IL). The sequencing results were analyzed and compared for sequence homology using the National Center for Biotechnology Information (NCBI) database and BLAST nucleotide tool, available at https://blast.ncbi.nlm.nih.gov.

### 2.2. Phylogenetic analysis

Sequencing of each amplified D1/D2 domain of the 26S rRNA gene was used to analyze the phylogenetic relationships among wild yeasts, using MEGA software (Molecular Evolutionary Genetics Analysis) v.10.1.7 (https://www.megasoftware.net) for alignment, construction and visualization of the phylogenetic tree. Sequence alignments were performed using ClustalW, which was also used to generate the phylogenetic trees using the bootstrap method with 1000 replications through neighbor-joining statistical method.

### 2.3. Fermentation tests

Isolated yeasts were inoculated into two tubes containing 5 mL YPD medium and incubated for 48 h at 30°C until obtaining a density of approximately 10^9^ CFU/mL. One of the cultures was used to inoculate 400 mL of 100% Pilsner malt extract (Briess) medium, density 1.040 g/cm^3^ and pH 5.4, while the second was used to inoculate 400 mL of India Pale Ale beer wort, 1.049 g/cm^3^, pH 5.5, 65 IBU, produced and supplied by Upland Brewering Company (Bloomington, IN, USA). Fermentation flasks were incubated without agitation, with an air-lock to release CO_2_, for 14 days at room temperature (approximately 23°C). During fermentation, whether visible fermentation activity (e.g., gas release through the air-lock) and the formation of a biofilm on the liquid surface was observed. After 14 days, samples were collected to analyze the final density and pH, allowing observation of acidification ability and apparent attenuation by each wild yeast isolated. Density was analyzed using a MISCO digital refractometer and pH using an Accumet AB150 (Thermo Fischer Scientific) pH meter.

### 2.4. Evaluation of important brewing characteristics of *M. megachiliensis* isolate ONP131

*M. megachiliensis* ONP131 and the commercial ale yeast *Saccharomyces cerevisiae* WLP001 (White Labs, San Diego, CA) were inoculated in 5 mL of YPD medium (pH 5.0), and incubated overnight with agitation at 30 °C. The optical density of both cultures was evaluated using a Beckman Coulter DU730 UV/Vis Spectrophotometer, and then cultures were diluted with ultrapure water to an OD_660nm_ of 0.06. Tests for the characterization of *M. megachiliensis* ONP131 were performed in 96-well plates, in duplicate, three independent times. For each well, 100 µL of the dilution of each yeast was mixed with 100 µL of the medium to be tested, at a concentration of 2x, and overlaid with 50 µL of mineral oil to avoid evaporation. Growth curves were followed over 48 h at 30°C by optical density readings every 15 min using a Synergy H1 Plate Reader (Biotek, Winooskim VT) with Gen5 Microplate Reader and Imager Software (Biotek, Winooski, VT). For all tests *S. cerevisiae* WLP001 was used as control yeast.

To evaluate important characteristics for use in beer production, different treatments were tested. Initially, the growth of *M. megachiliensis* was evaluated in standard YPD medium, as well as in dry malt extract (DME), with densities of 1.020 g/cm^3^ and 1.040 g/cm^3^, respectively. Cultures in YPD were subjected to different incubation temperatures at 30°C, 34°C, and 37°C, seeking to assess tolerance to higher temperatures for growth. Consumption of different carbohydrates was tested in the presence of 2% (w/v) glucose, 2% maltose, 2% sucrose, or 2% galactose, while tolerance to osmotic stress was assessed using concentrations of 10, 20 and 30% (w/v) glucose and maltose. All of the above conditions were analyzed with YP as a basis for medium formulation (10 g/L yeast extract and 20 g/L peptone). Tolerance to salinity was tested using YPD + 0.5, 1, 5, or 10% (w/v) NaCl. Ethanol tolerance was evaluated in YPD medium supplemented with 2, 4, 5, or 6% (v/v) ethanol. As a control, tests were performed using only YPD to evaluate the growth kinetics and final OD_660nm_ of *M. megachiliensis* and *S. cerevisiae* cultures.

The ability to grow at different pHs was evaluated by incubating *M. megachiliensis* in YPD medium at pH = 2.5, 3.0, 3.5 or 5.0 (adjusted with 1 M NaOH). To observe the impact of lactic acid on yeast cultivation, YPD medium was adjusted with 85% lactic acid (to a final pH of 2.5, 3.0, 3.5, or 5.0 prior to yeast inoculation). The ability to tolerate different concentrations of iso-α-acids (compounds present in hops that have antimicrobial activity) was also evaluated in YPD medium, with concentrations ranging from 10 to 200 ppm of 30% isomerized hop extract (Hopsteiner, Mainburg, DE). On YPD agar plates supplemented with (a) 30% isomerized hop extract at concentrations of 90 and 120 IBU (International Bitterness Units) or (b) β-acids, from 45% Beta Bio (Hopsteiner, Mainburg, DE) at concentrations of 100 ppm and 200 ppm, spot test were performed using 5 µL of 10^8^-10^3^ CFU/mL culture dilutions to confirm tolerance to hop compounds.

### 2.5. Statistical analyses

All statistical analyses were performed using GraphPad Prism 7 software. Differences between values were compared using analysis of variance (ANOVA) and Tukey test, where *p*-values < 0.05 were considered statistically significant.

## 3. Results

### 3.1. Wild yeasts from Olympic National Park samples

Sixteen yeast isolates were obtained from fifteen samples collected in Olympic National Park. All isolates were subjected to fermentation tests in malt extract and beer wort. Yeasts displaying the most promising fermentative potential were identified by PCR, and we identified six strains from four different species: *Debaryomyces hansenii* (ONP21, GenBank accession n. MZ506604; ONP27, GenBank accession n. MZ506605), *Yamadazyma scolity* (ONP63, GenBank accession n. MZ506606), *M. megachiliensis* (ONP93, GenBank accession n. MZ506607; ONP131, GenBank accession n. MZ506609), *Starmerella riodocensis* (ONP96, GenBank accession n. MZ506608). We observed that the colony morphologies of the *D. hanseniii, S. riodocensis* and *Y. scolity* isolates on WLN and YPD plates were similar to *S. cerevisiae* WLP001, being circular with smooth margins, though they did not show elevation (Supplementary Information – Fig. S1). In contrast, *M. megachiliensis* ONP93 and ONP131 formed colonies with filamentous borders, easily distinguishable from the other yeasts for having an olivaceous black color after 72 h of growth. Microscopic analysis showed cells with almost twice the size of *S. cerevisiae*, that formed clumps. After 24 h of aerated cultivation in liquid YPD medium, we also observed that *M. megachiliensis* isolates sedimented as popcorn-like clumps after 10 s without agitation (Fig. S2).

Based on the D1/D2 sequences of the 26S rRNA genes of the above yeasts, a phylogenetic tree was constructed to determine relationships among the isolates (Fig. 1). The yeasts *D. hansenii* and *Y. scolity* are taxonomically classified in the same family, Debaryomycetaceae (which explains their proximity in phylogeny), and compared to *S. cerevisiae* and *S. riodocensis*, they all belong to same class of Saccharomycetes. This class comprises those considered “true yeasts”, which share ultrastructural characteristics such as aspects of nuclear division and ascospore formation (Blackwell and Spatafora, 2004). All of the abovementioned yeasts are also part of the Ascomycota phylum, while *M. megachiliensis* is classified in the phylum Basidiomycota, which explains its phylogenetic distance from the others.

**Fig. 1:**
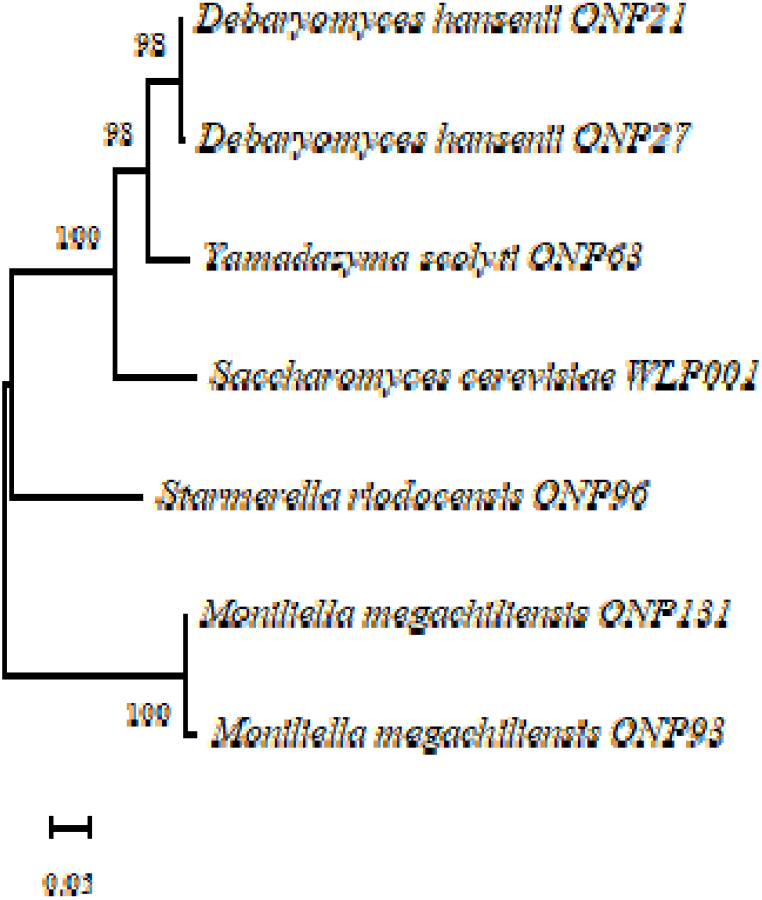
Phylogenetic tree of wild yeasts identified from Olympic National Park samples and a commercially available *S. cerevisiae* yeast. The D1/D2 rRNA sequences of the isolated strains were aligned, and the phylogenetic relationships among them were constructed using the bootstrap method with 1000 replicates, based on neighbor-joining analysis.

### 3.2. Fermentation tests with the wild yeasts isolates

Fermentation tests results are shown in Table 1. In malt extract and beer wort fermentations by *D. hansenii* ONP21 and ONP27, we observed that, in addition to the final pH and density being higher than ideal for beers, production of non-pleasant aromas that are undesirable for fermented beverages were detected. Although off-flavors were not noticed in *Y. scolity* ONP63 and *S. riodocensis* ONP96 fermentations, their inability to adapt to beer wort suggests that these isolates are not suitable for application in this fermentation process. *M. megachiliensis* isolates ONP93 and ONP131 fermentations displayed final pH, density, and alcohol by volume (ABV) values closer to those found in fermentations by *S. cerevisiae* WLP001, in addition to presenting interesting flavors similar to those detected in beers with esterified aromas.

**Table 1:**
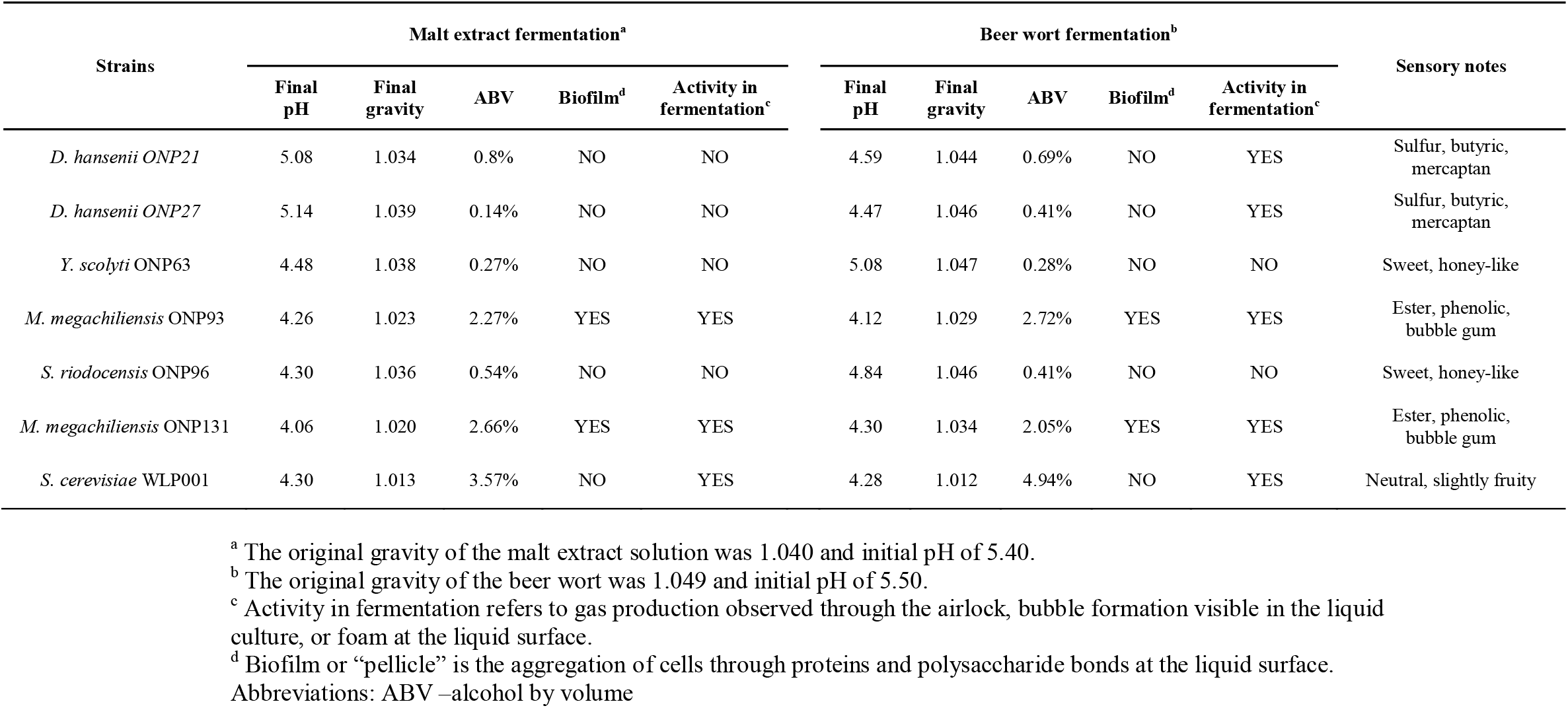
Fermentation tests of malt extract and beer wort using wild yeasts isolated from Olympic National Park. Preliminary sensory notes were collected as described previously (Osburn, Ahmad, and Bochman 2016).

In Figure 2, we show that *M. megachiliensis* isolates displayed an apparent attenuation of carbohydrates present in malt extract and beer wort between 30 and 50%, demonstrating a 3 to 4-fold greater capacity to attenuate wort compared to the other isolates. Because the *M. megachiliensis* ONP131 isolate performed slightly better than ONP93 and was informally ranked as producing better beer, it was selected for subsequent tests and physiological characterization.

**Fig. 2:**
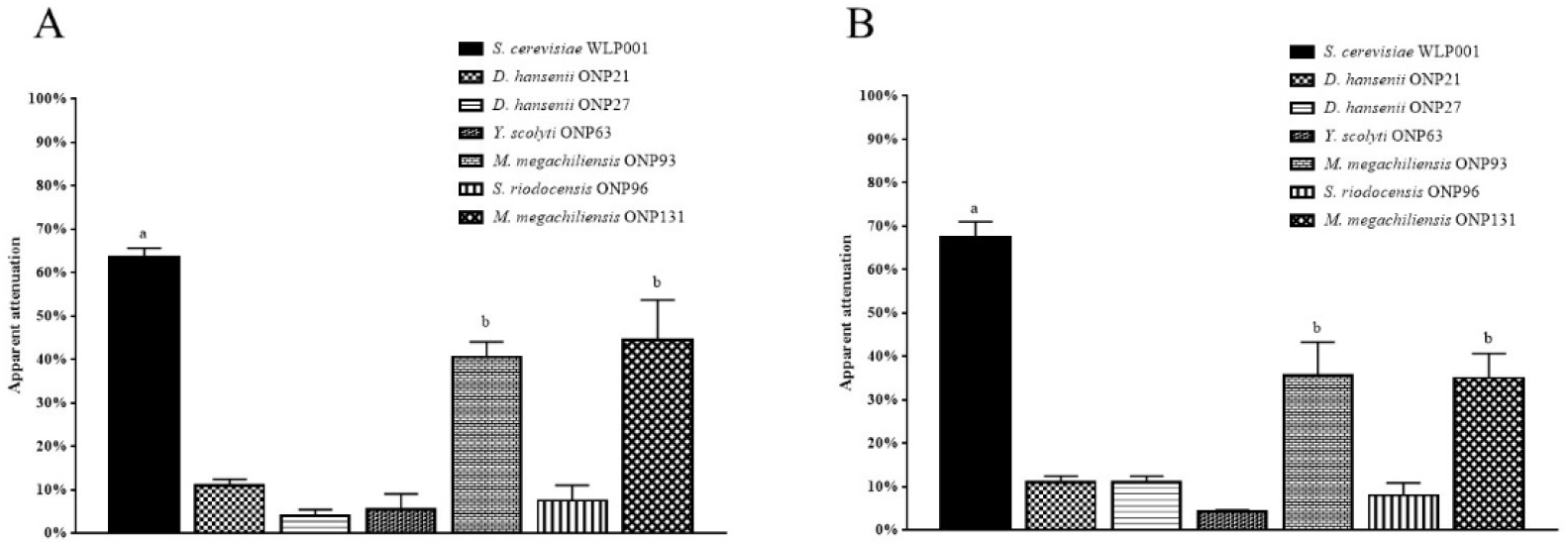
Apparent attenuation in malt extract (A) and beer wort (B) by the isolated yeasts. Commercial available *S. cerevisiae* was used as a control for normal attenuation by brewer’s yeast. Error bars indicate ± SD. Letters indicate statistical difference (p < 0.05): **a**) difference from all yeasts; **b**) different from all yeasts, except for the same yeast species.

### 3.3. Characterization of *M. megachiliensis* ONP131 brewing characteristics

The growth capacity of *M. megachiliensis* isolate ONP131 was evaluated under different conditions and compared to a standard ale strain of *S. cerevisiae*. Growth curves are presented in Figure S3, while the following results are related to the final OD_660nm_ measured in *M. megachiliensis* ONP131 and *S. cerevisiae* WLP001 cultures.

No significant differences were observed in the absorbance of the two yeasts when cultivated in YPD medium (*p* < 0.05) (Fig. 3A), but there was a tendency of lower growth when *M. megachiliensis* was cultivated in DME with densities of 1.020 and 1.040, showing lower final absorbances in both cases. The incubation temperature directly influenced the capacity of *M. megachiliensis* to flourish, demonstrating a decrease of approximately 40% in final absorbance of cultures at 34°C and 37 °C relative to 30°C (Fig. 3B).

**Fig. 3:**
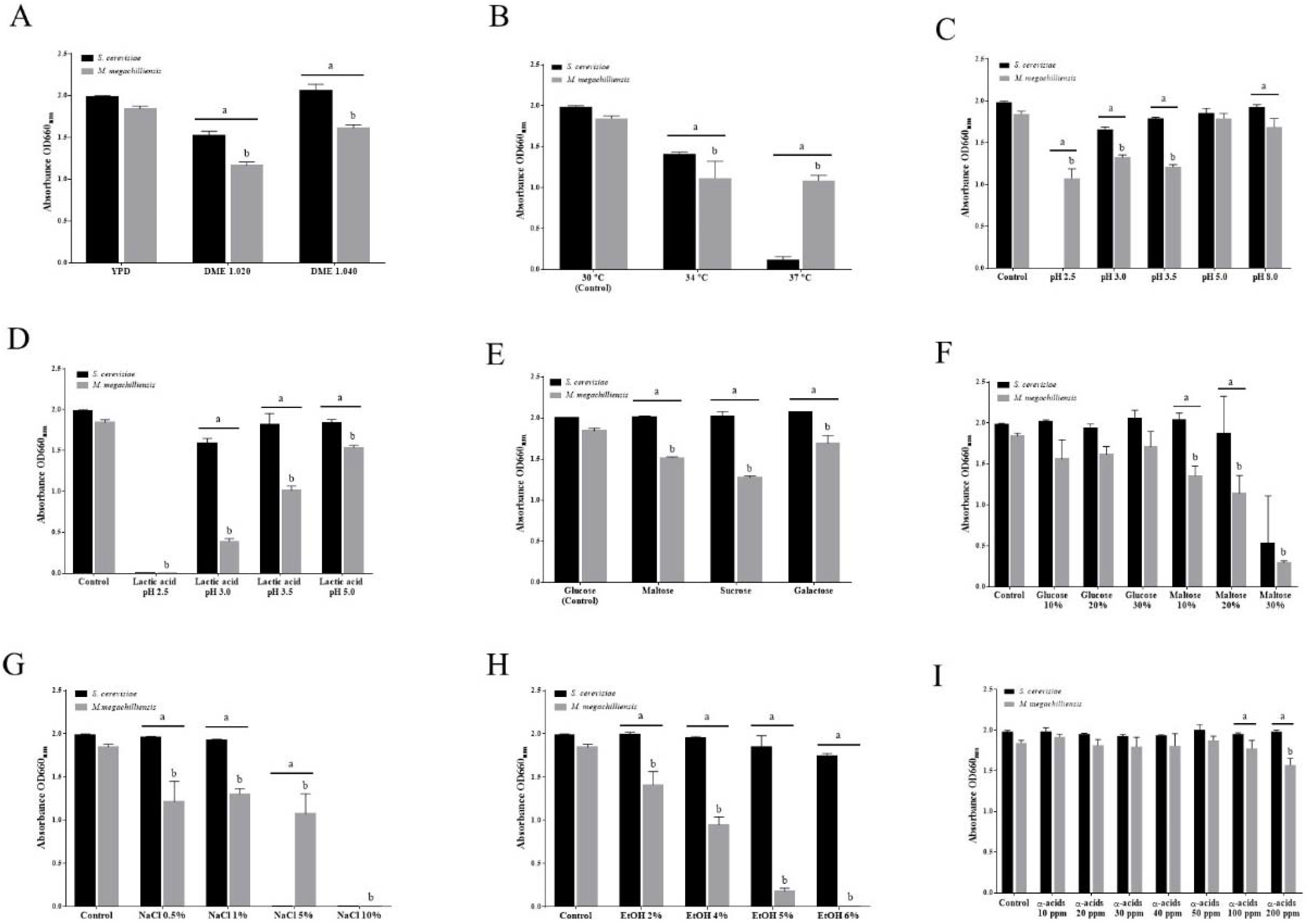
*M. megachiliensis* physiological characterization for beer fermentation. Characteristics of *M. megachiliensis* ONP131 regarding growth medium for yeast propagation (A), incubation temperature (B), optimum culture pH (C), tolerance to lactic acid (D), growth ability in media containing other carbohydrates as a carbon source (E), osmotic tolerance to glucose and maltose (10% > 30%) (F), halotolerance in NaCl-supplemented media (G), ethanol tolerance (H), and resistance to α-acids from hops extract (I). Data are expressed as the terminal O.D._660nm_ of the cultures. Error bars indicate ± SD. Letters indicate statistical differences (*p* < 0.05): **a**) difference between the growth of *S. cerevisiae* WLP001 (control yeast) and *M. megachiliensis* cultures; **b**) differences between the control group and analyzed value.

Medium acidification with HCl to pH 2.5 hindered the growth of both *S. cerevisiae* and *M. megachiliensis*, but ONP131 maintained an OD_660nm_ ≥ 1.0, representing at least 50% of its original growth observed in control culture (pH 6.0) (Fig. 3C). Although a significant (*p* < 0.05) decrease in growth was observed in the range of pH 2.5-3.5, cultures subjected to pH 5.0 and 8.0 displayed growth equivalent to that of the control. Acidifying the culture medium with lactic acid proved to be more harmful to yeast growth, preventing it at pH 2.5 or limiting it to an absorbance ≤ 1.0. Even the medium with low amounts of added lactic acid (pH 5.0) was also harmful to the growth of both yeasts (Fig. 3D).

While comparing the growth of *M. megachiliensis* and S. cerevisiae, we next assessed the effects of various sugars. In general, other carbohydrate sources are not metabolized by yeast as effectively as glucose. When applied at a concentration of 2%, maltose, sucrose and galactose resulted in cultures with a lower absorbance (OD_660nm_ = 1.51, 1.27, and 1.68, respectively) than that detected for the control medium containing glucose (OD_660nm_ = 1.84) and for *S. cerevisiae* (Fig. 3E). Osmotic pressure tolerance analysis using glucose and maltose revealed that *M. megachiliensis* tolerates glucose concentrations up to 30%, having no significant differences in final optical density (*p* < 0.05) (Fig. 3F). However, concentrations above 10% maltose caused changes in growth, with 30% maltose significantly limiting growth to OD_660nm_= 0.3, which is only 15% of that observed in the control YPD medium.

Halotolerance is a characteristic of some yeasts, and this ability to tolerate high salt concentrations was also evaluated in our study. We found that *M. megachiliensis* tolerated up to 5% NaCl, while the same concentration greatly limited the growth of *S. cerevisiae* (Fig. 3G). Supplementation of the culture medium with ethanol next revealed that ONP131 growth decreased as ethanol concentration increased up to 6% ethanol, which completely prevented *M. megachiliensis* growth. (Fig. 3H). Among limiting conditions for growth in beer wort, the α-acids derived from hops are known to inhibit bacterial growth but can also affect yeasts at high concentrations {Hazelwood, 2010 #5002}. In our study we observed that only very high concentrations (≥ 200 ppm) of α-acids were able to impact *M. megachiliensis* growth (Fig. 3I); below that, no statistical differences were observed compared to control cultures. Tolerance was also observed for α-acids (90 and 120 IBU) and β-acids (100 ppm and 200 ppm) used as supplements in solid culture media, in which *M. megachiliensis* showed similar growth to *S. cerevisiae*, demonstrating resistance to these conditions (Fig. 4).

**Fig. 4:**
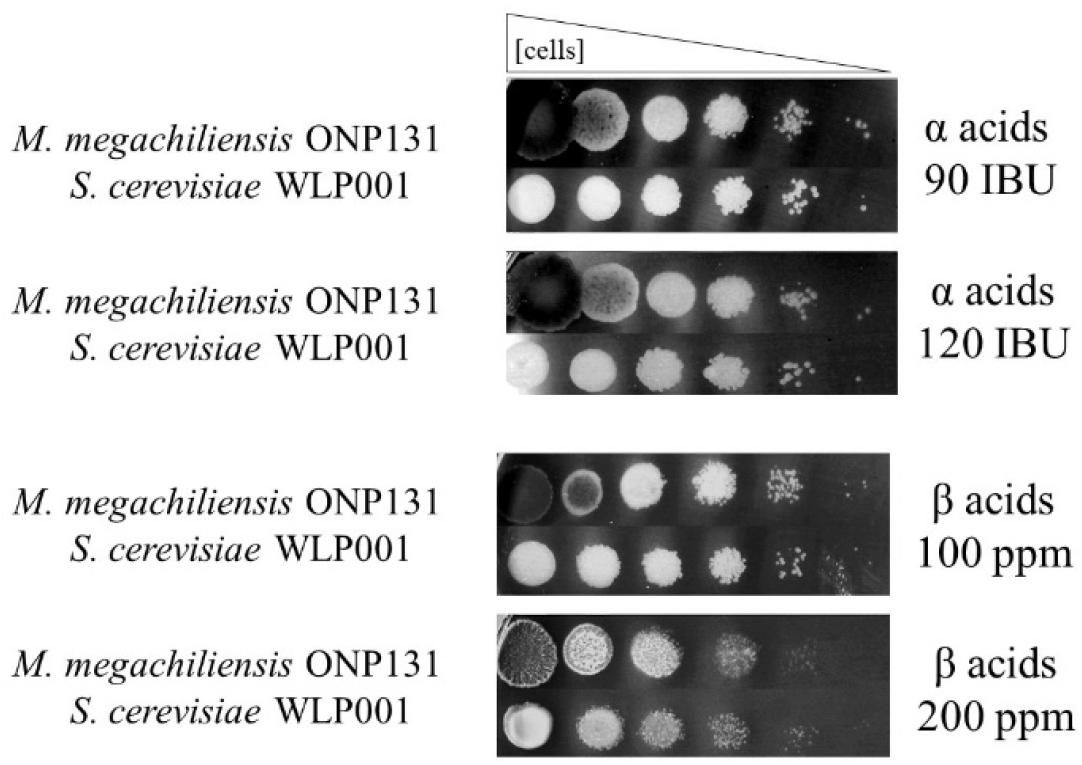
*M. megachiliensis* tolerance to α- and β-acids from hops. *M. megachiliensis* ONP131 and *S. cerevisiae* WLP001 (control) were cultured on YPD plates supplemented with high concentrations (90 or 120 IBU) of iso-α-acids (30% isomerized hop extract) or (100 ppm and 200 ppm) β-acids (45% Beta Bio). Both yeasts showed similar growth under these conditions, confirming *M. megachiliensis* tolerance to the antimicrobial components present in hops.

## 4. Discussion

When bio-prospecting, yeasts are generally found associated with bees and flowers (Lachance et al., 2001; Pimentel et al., 2005) as well as from a wide range of other sources, such as processed food products, soil, marine environments, and fermented products (Suzuki et al., 2011). Here, we isolated novel strains from small plant roots, tree bark and flowers, which are sources similar to those described elsewhere in the literature. Of the six strains displaying some fermentative potential, low metabolism of wort sugars was observed for the ONP21 and ONP27 isolates of *D. hansenii*, yielding apparent attenuation between 5 and 10% in both tests. Used in many processes such as the fermentation of cacao and meat products (Ramos et al., 2017), *D. hansenii* is known to have great biotechnological potential. Concerning its application in the brewing industry, some strains have interesting features, such as conversion of ferulic acid to 4-vinyl-guaiacol (Yaguchi et al., 2017). However, production of unpleasant aromas by ONP21 and ONP27 such as sulfur, butyric acid, and mercaptans was easily detectable by simple sensory analysis in our study, which was considered a problem for their future application in beer fermentation.

The inability of *S. riodocensis* ONP96 to utilize maltose for its development (Pimentel et al., 2005) was detected in both fermentation tests, resulting in an apparent attenuation of 5-10%. *Y. scolity* has been reported to be able to grow using maltose, however the apparent attenuation observed suggested it was unable to ferment this carbohydrate in our study (apparent attenuation between 3 and 10%) and others (Kurtzman, 2011). Our results also indicated that neither *Y. scolity* ONP63 nor *S. riodocensis* ONP96 contributed to the aroma *bouquet* of fermented malt extract or beer wort, suggesting that these strains have no potential for application in the fermentation of beers even when co-inoculated with strains better able to metabolize maltose. This left the *M. megachiliensis* isolates to test further.

Despite the fact that the relationship between the classes of fungi within the Basidiomycota phylum is controversial, the *Moniliella* genus represents yeast-like fungi that can be identified by ellipsoidal budding cells that form terminally on true hyphae, pseudomycelium, or chlamydospores and which are capable of fermenting various sugars, including maltose (Blackwell and Spatafora, 2004; Hoog et al., 2011; Rosa et al., 2009). As for this writing, there are no reports in the literature related to the application of *M. megachiliensis* in fermented beverage production, so it was necessary to characterize its fermentation capacity, as well as the flavors and aromas originating from fermentation. Although they do not have an attenuation profile like *S. cerevisiae*, it is known that yeasts with a low or medium attenuation percentage can have a strong contribution to the flavors of a beer (Michel et al., 2016). The esterified and bubble gum aromas observed in our study are in agreement with the work by Hoog et al. (2011) and Thanh and Hien, (2019), in which they report “fruity” and “peach-like” aromas derived from ethyl-acetate, γ-decalactone, and acetaldehyde in media cultivated with yeasts of the *Moniliella* genus. Thus, because it was responsible for the production of pleasant aromas and had moderate attenuation capacity, our *M. megachiliensis* ONP131 isolate was selected for further study.

There is a need to know the performance of wild yeasts through their domestication to establish the feasibility of using these organisms for brewing purposes (Steensels and Verstrepen, 2014). As described by Rakete and Glomb (2013), 90% of beer wort is composed of glucose, sucrose, maltose, and maltotriose, usually with a higher concentration of maltose. Therefore, tests were performed to characterize yeast growth using mainly the carbohydrates present in wort, in which we observed the ability of *M. megachiliensis* to grow in media where these carbohydrates were used as the sole carbon source. The possibility of cultivating yeast in DME also allows homebrewers and craft breweries to propagate this isolate in-house and maintain successive controlled fermentations.

Thermotolerance in yeasts is an uncommon feature, but those able to withstand temperatures ≥ 37°C are of interest for industrial use because they require lower costs for bioprocesses cooling and reduce contamination risks in these systems (Lehnen et al., 2019). We found that *M. megachiliensis* was able to sustain its growth when incubated in the 34-37°C range, even though the final absorbance value was lower than that observed for growth at 30°C. Therefore, this yeast has potential for application in bioprocesses that demand higher temperatures, as has already been demonstrated for the thermotolerant yeasts *Kluyveromyces marxianus* and *Ogataea polymorpha* (Lehnen et al., 2019).

Increases in osmotic pressure cause changes in the viability, size, and shape of lager and ale yeasts, which influence their fermentation capacity (Pratt et al., 2003). Osmotolerance is strain-dependent, being derived from membrane structure, vacuolar function, residual levels of trehalose, and especially the abundance of osmoprotective macromolecules (Gibson et al., 2007). In media containing 10, 20 or 30% glucose, no statistical difference was observed in the final absorbance of the *M. megachiliensis* cultures. However, when the same maltose concentrations were analyzed, it was observed that there was a decrease in OD_660nm_ due to high osmotic pressure and inability to metabolize this carbohydrate. Although different enzymatic pathways are activated in response to high extracellular osmolarity (e.g., the high-osmolarity glycerol pathway), at certain concentrations, these are not sufficient to ensure cell development, consequently culminating in a decrease of cell viability and concentration (Gibson et al., 2007).

Exploring the ability of yeasts to tolerate salt concentrations (ionic stress) is important in industrial fermentations, where salt can favor yeast growth, enhance ethanol production, and at the same time, reduce the risk of contamination by microorganisms with low halotolerance (Corte et al., 2006). While *S. cerevisiae* was not able to grow in 5% NaCl, *M. megachiliensis* sustained its growth and displayed halotolerance up to 5% NaCl with no significant difference in absorbance compared to growth with at 0.5% and 1% NaCl. At 10% NaCl concentration, however, the yeast was not able to survive.

Ethanol inhibits cell growth, viability, and fermentation rate in yeast (Pratt et al., 2003). Yeast ethanol tolerance can be assessed by exposing the yeast culture to ethanol at different concentrations until cellular growth suppression occurs, an important tool to characterize yeast species and strains considered for application in alcoholic fermentations (Da Silva et al., 2013). We observed that the growth of *M. megachiliensis* decreased with increasing ethanol concentration in the medium, which is similar to the pattern displayed by other wild yeast isolates characterized by (Lee et al., 2011). Ethanol at a 6% concentration was identified as a limiting factor for *M. megachiliensis* ONP131 growth, which has similarly been described as a limiting factor for several other wild yeasts, such as *Hanseniaspora uvarum, Lachancea kluyveri*, and *Torulaspora delbrueckii* (Osburn et al., 2016). Therefore, it is suggested that *M. megachiliensis* is unable to survive in fermentations that exceed this ethanol concentration and should be used to produce beer below 5% ethanol, though adaptation to increasing ethanol concentration during fermentation (rather than immediate exposure to 6% ethanol upon inoculation) may enable *M. megachiliensis* ONP131 to survive in higher ABV beers.

It is well known that pH is another limiting factor for the proper growth of microorganisms, as well as the kinetics of pH decrease during fermentations by some microorganisms (Narayanan et al., 2016). Even though yeast cells are able to maintain appropriate internal pH by utilizing cell buffer systems and other pathways during changes in external pH conditions, certain pH ranges are not tolerated by yeasts (Brandão et al., 2014). In our tests, we found that *M. megachiliensis* was able to maintain its growth in YPD medium adjusted to pH 2.5 with HCl, but when the pH was adjusted to 2.5 with lactic acid, growth was not observed. The acidic shock occasioned by high levels of lactic acid was also observed by Rogers et al. (2016) for *S. cerevisiae*, which generally fail to carbonate sour beers with low pHs. The ability to withstand a pH range ≥ 3.0 (in YPD adjusted with HCl or lactic acid) makes to application of *M. megachiliensis* ONP131 in sour beers (pH ∼3.4) interesting, whether for complete wort fermentation or post-fermentation, during the natural carbonation process.

Due to large concentration of hops in certain beer styles (e.g., India Pale Ales), we were also interested in determining the capacity of *M. megachiliensis* to tolerate compounds present in hops, such as α- and β-acids. As demonstrated by Methner et al. (2019) and Michel et al. (2016), the resulting inhibitory effect of these compounds may vary according to the yeast species analyzed, of which not all wild isolates are able to grow in concentrations up to 100 IBU and 200 ppm of β-acids. Here, though, we found that *M. megachiliensis* was able to maintain its growth at levels close to that observed for the control (no α-acids added), as well as able to tolerate β-acids concentrations between 100-200 ppm. Thus, we identified hop tolerance by the ONP131 isolate as evidence that it can be used in the production of highly hopped beers.

## 5. Conclusion

In this work, we successfully isolated different yeast species from environmental samples collected in Olympic National Park. Upon analysis of fermentative activity by these yeast isolates, we selected *M. megachiliensis* ONP131 for further physiological characterization, of its growth in conditions generally found in beer worts. Overall, we found that *M. megachiliensis* ONP131 is an isolate with potential for the brewing industry, perhaps when co-inoculated with a typical *Saccharomyces* beer strain to overcome the low attenuation of ONP131. Based on its halo- and thermotolerance, *M. megachiliensis* ONP131 may also be of interest for other bioprocesses as well.

