## Supplementary Information for "Isolation of wild yeasts from Olympic National Park and *Moniliella megachiliensis* ONP131 physiological characterization for beer fermentation"

**Title:**

¹Laboratório de Microbiologia, Centro de Desenvolvimento Tecnológico, Universidade Federal de pelotas, Pelotas, RS, Brazil

²Bochman Lab, Molecular and Cellular Biochemistry Department, Indiana University Bloomington, Bloomington, IN, United States

**Corresponding author:**

Prof. PhD. Matthew L. Bochman,

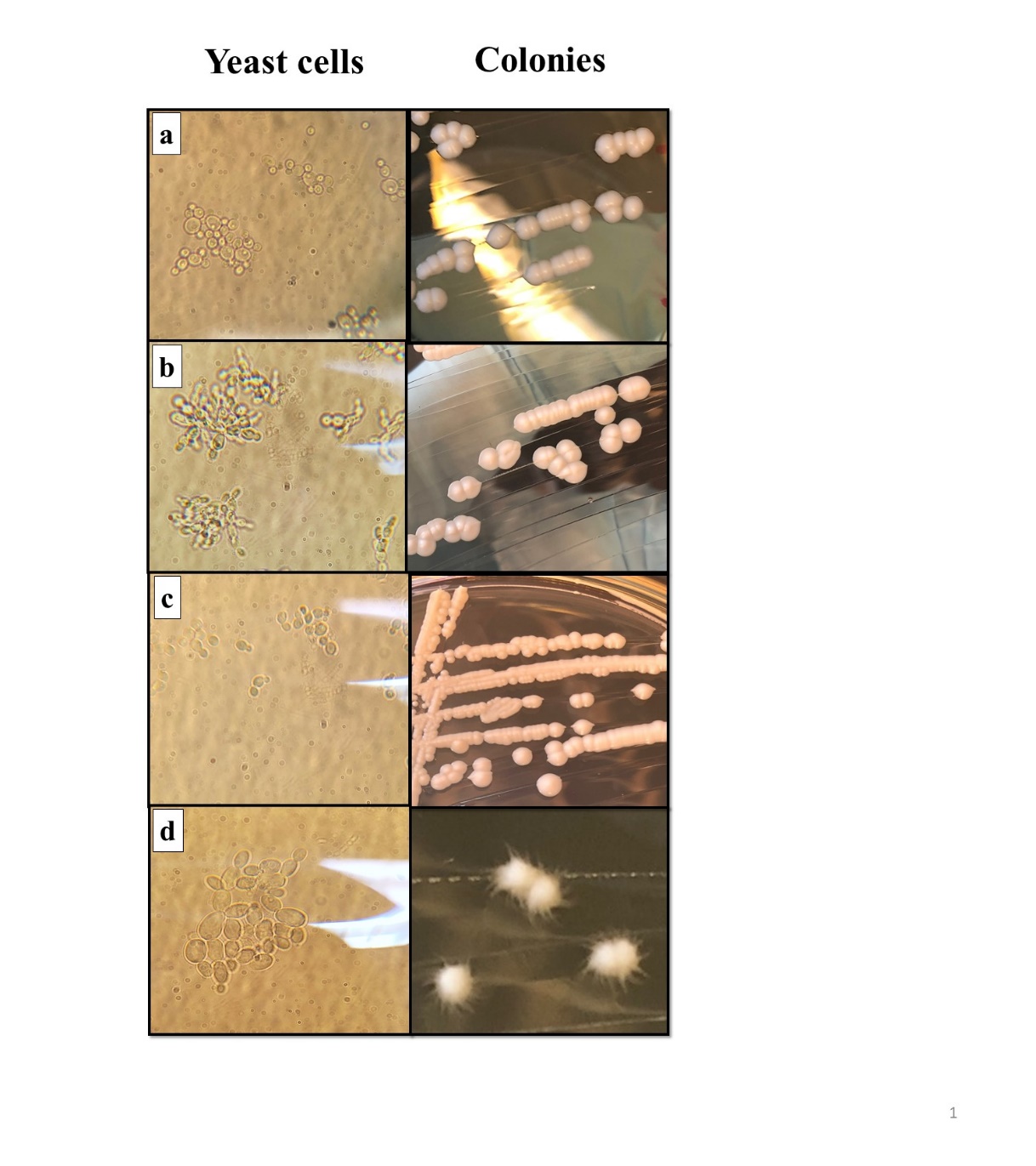
**Fig. S1:** Micrographs of isolated yeasts, taken under at x1000 magnification (left), and their respective colony morphologies (right), after 48 h on YPD agar at 30°C**.** a) *D. hansenii*, b) *Y. scolyti,* c) *S. riodocensis* and d) *M. megachiliensis*.

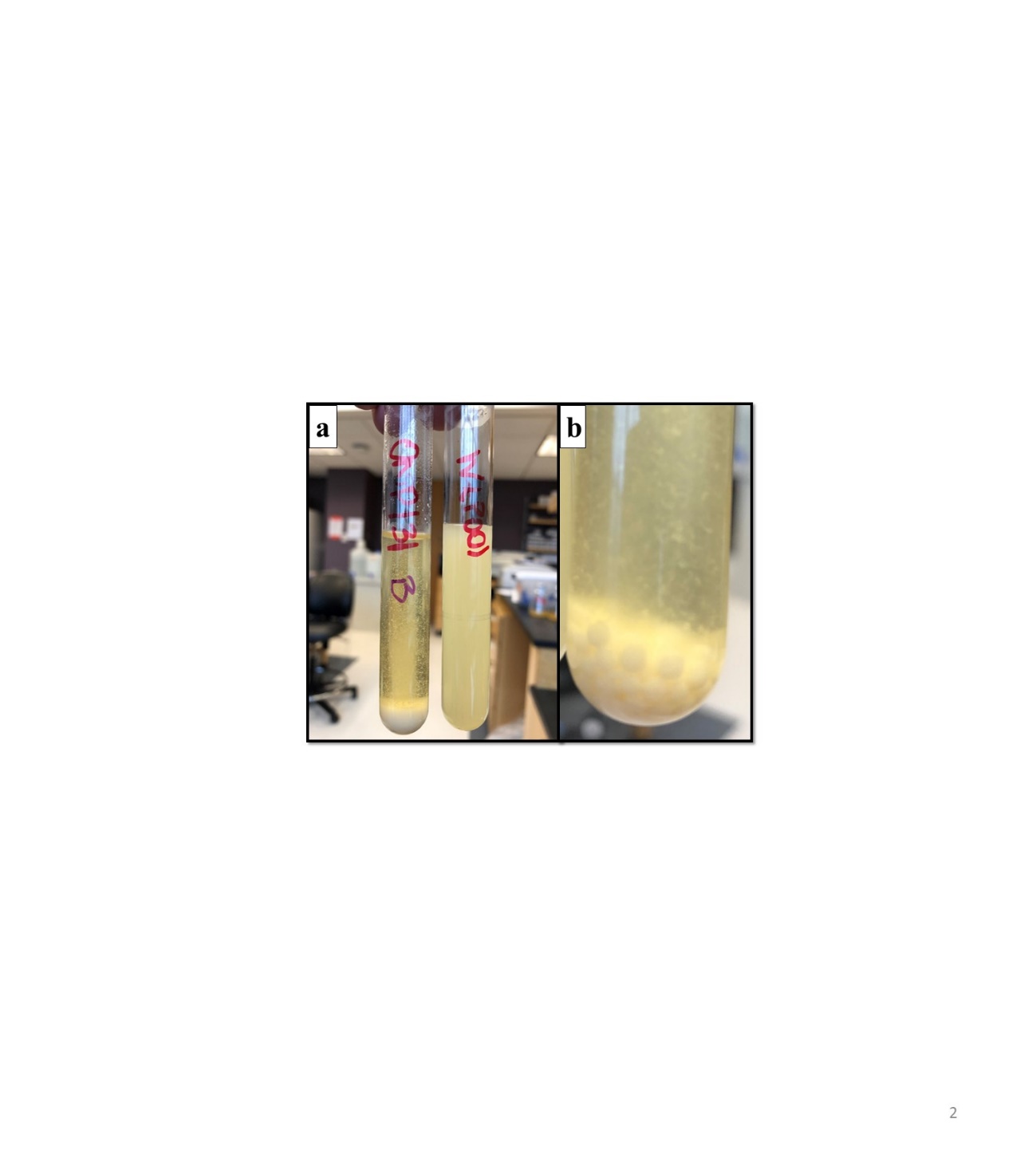
**Fig. S2: Sedimentation of *M. megachiliensis* cultures after 10 s with no shaking.** a) Yeast cultures after 24 h in 10 mL of YPD. The tube on the right is a *S. cerevisiae* WLP001 control culture. b) Popcorn-like clumps of sedimented ONP131 cells.

**Fig. S3: Growth curves of *M. megachiliensis* ONP131 cultivated under different conditions.**
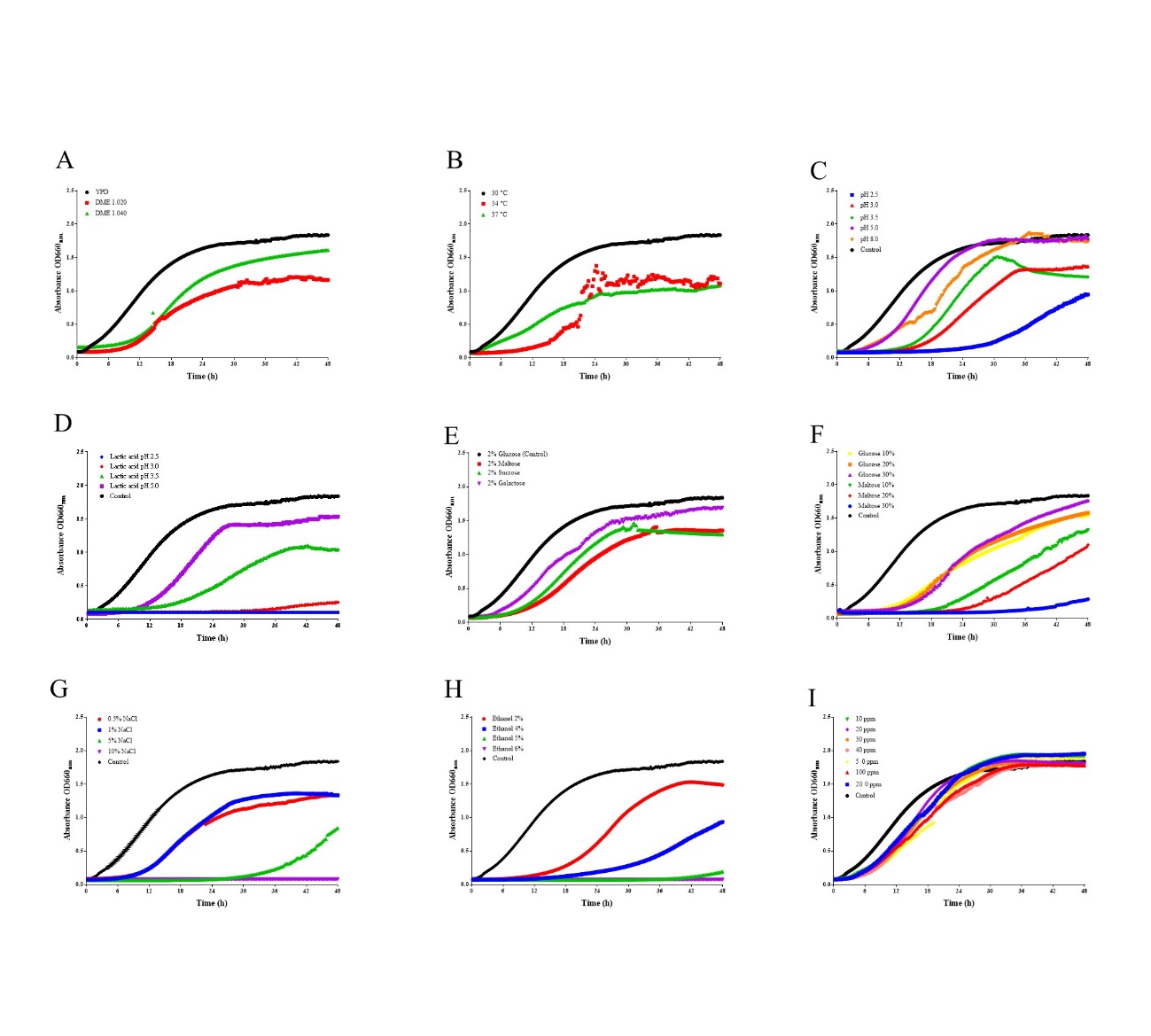
Medium for yeast propagation (A), incubation temperature (B), optimum culture pH (C), tolerance to lactic acid (D), growth in media containing other carbohydrates as a carbon source (E), osmotic tolerance to glucose and maltose (10 to 30%) (F), halotolerance in NaCl-supplemented media (G), ethanol tolerance (H), and resistance to α-acids from hops extract (I). Growth curves were followed over 48 h at 30°C by optical density readings every 15 min on a Synergy H1 Plate Reader (Biotek, Winooskim VT) with Gen5 Microplate Reader and Imager Software (Biotek, Winooski, VT). Averaged data are expressed in absorbance (OD_660nm_) values by time. Error bars representing standard deviations were omitted for data clarity.
